## supplementary information for "Enalapril mitigates senescence and aging-related phenotypes by targeting antioxidative genes via phosphorylated Smad1/5/9"

**Supporting Information**

#### Supplementary Tables

**Table S1 Primers used in RT-qPCR**

| Name | Primer sequence |
|---|---|
| IL1β | For: ATGATGGCTTATTACAGTGGCAA<br>Rev: GTCGGAGATTCGTAGCTGGA |
| IL6 | For: ACTCACCTCTTCAGAACGAATTG<br>Rev: CCATCTTTGGAAGGTTCAGGTTG |
| CXCL10 | For: GTGGCATTCAAGGAGTACCTC<br>Rev: TGATGGCCTTCGATTCTGGATT |
| CXCL16 | For: CCCGCCATCGGTTCAGTTC<br>Rev: CCCCGAGTAAGCATGTCCAC |
| CCL2 | For: AAGACCATTGTGGCCAAGGA<br>Rev: TTCGGAGTTTGGGTTTGCT |
| MMP2 | For: CCCACTGCGGTTTTCTCGAAT<br>Rev: CAAAGGGGTATCCATCGCCAT |
| BMPR1A | For: CTTTACCACTGAAGAAGCCAGCT<br>Rev: AGAGCTGAGTCCAGGAACCTGT |
| ID1 | For: CTGCTCTACGACATGAACGG<br>Rev: GAAGGTCCCTGATGTAGTCGAT |
| ID2 | For: AGTCCCGTGAGGTCCGTTAG<br>Rev: AGTCGTTCATGTTGTATAGCAGG |
| ID3 | For: GAGAGGCACTCAGCTTAGCC<br>Rev: TCCTTTTGTCGTTGGAGATGAC |
| ACTB | For: CACCCCGTGCTGCTGAC<br>Rev: CCAGAGGCGTACAGGGATAG |
| TXN | For: GTGAAGCAGATCGAGAGCAAG<br>Rev: CGTGGCTGAGAAGTCAACTACTA |
| GPX4 | For: GAGGCAAGACCGAAGTAAACTAC<br>Rev: CCGAACTGGTTACACGGGAA |
| PRDX5 | For: CTTCACCCCTGGATGTTCCAA<br>Rev: AGGCATCATTAACACTCAGACAG |

22 **Table S2 Primers used in pSmad1/5/9 ChIP-qPCR**

| Name | Primer sequence | Notes |
|---|---|---|
| HPRT1 | For: TGTTTGGGCTATTTACTAGTTG<br>Rev: ATAAAATGACTTAAGCCCAGAG | Negative control<br>(Morikawa et al., 2011;<br>Sullivan & Santos, 2020) |
| HBB | For: GGGCTGAGGGTTTGAAGTCC<br>Rev: CATGGTGTCTGTTTGAGGTTGC | Negative control<br>(Morikawa et al., 2011) |
| ID1 | For: AGTCCGTCCGGGTTTTATG<br>Rev: TGTGTCAGCGTCTGAACCAG | Positive control<br>(Morikawa et al., 2011;<br>Sullivan & Santos, 2020) |
| ID2 | For: ACTCTATTTACCACCCCAGC<br>Rev: AGCTTCCCTTCGTCCCCAT | |
| TXN | For: CAGGGCTGGATTCCTCGAAA<br>Rev: CAAGGACGTACACACCGAGA | |
| PRDX5 | For: GTATGGGACTAGCTGGCGTG<br>Rev: TCACTGTACCGTCTTGCTGC | |
| GPX4 | For: GAAGCAGAGACGGGAGGTTC<br>Rev: CTTGTGTCTAGGAGGCCGTG | |
| SOD3 | For: TCTGAGGGGTTAGTGGGGAG<br>Rev: CCCCTTTCTCGTCACTCCAG | |

23 **Table S3 shRNA target sequence used in knock down experiment**

| Name | Target Sequence |
|---|---|
| shID1 #1 | CCGGCCTACTAGTCACCAGAGACTTCTCG<br>AGAAGTCTCTGGTGACTAGTAGGTTTTT |
| shID1 #2 | CCGGACTCGGAATCCGAAGTTGGAACTC<br>GAGTTCCAACTTCGGATTCCGAGTTTTTT |
| shID2 #1 | CCGGCCCTTCTGAGTTAATGTCAAACTCG<br>AGTTTGACATTAACTCAGAAGGGTTTTT |
| shID2 #2 | CCGGCCCACTATTGTCAGCCTGCATCTCG<br>AGATGCAGGCTGACAATAGTGGGTTTTT |
| shBMPR1A | CCGGCGCCAATCTCATACAAGCCATCTC<br>GAGATGGCTTGTATGAGATTGGCGTTTTT |

**Table S4 Antibodies**

| Name | Source | Number |
|---|---|---|
| Mouse monoclonal anti-β-actin | Santa Cruz Biotechnology | Cat#sc-47778 |
| Rabbit monoclonal anti-CDKN2A/p16INK4a | Abcam | Cat#ab108349 |
| Rabbit monoclonal anti-p21 | Abcam | Cat#ab109199 |
| Rabbit monoclonal anti-Phospho-SMAD1 (Ser463/465)/ SMAD5 (Ser463/465)/ SMAD9 (Ser465/467) | Cell Signaling Technology | Cat#13820S |
| Rabbit monoclonal anti-SMAD1 + SMAD5 + SMAD9 (phosphor S463 + S465 + S467) | Abcam | Cat#ab92698 |
| Rabbit monoclonal anti-Phospho-SMAD5 (Ser463/465) | Huabio | Cat#ET1605-5 |
| Rabbit polyclonal anti-SMAD1/5/9 | Immunoway | Cat#YT4325 |
| Rabbit monoclonal anti-SMAD5 | Huabio | Cat#ET1606-26 |
| Rabbit monoclonal anti-SMAD4 | Abcam | Cat#ab40759 |
| Rabbit polyclonal anti-SMAD2 (Ser465/467) | Cell Signaling Technology | Cat#3101S |
| Rabbit monoclonal anti-SMAD3 (Ser423/425) | Cell Signaling Technology | Cat#9520S |
| Rabbit polyclonal anti-SMAD2/3 | Cell Signaling Technology | Cat#3102S |
| Rabbit polyclonal anti-BMP2 | Huabio | Cat#ER80602 |
| Mouse monoclonal anti-BMP4 | Huabio | Cat#EM1706-23 |
| Rabbit polyclonal anti-ID1 | Immunoway | Cat#YN0093 |
| Rabbit polyclonal anti-ID2 | Immunoway | Cat#YN2457 |
| Rabbit polyclonal anti-PRDX5 | ABclonal | Cat#A1269 |
| Rabbit monoclonal anti-TXN | ABclonal | Cat#A4024 |
| Rabbit monoclonal anti-TXN2 | ABclonal | Cat#A4424 |
| Rabbit polyclonal anti-NDUFB10 | ABclonal | Cat#A9382 |
| Rabbit monoclonal anti-GPX4 | Abmart | Cat#T56959 |
| IRDy 800CW Goat anti-Mouse IgG Secondary Antibody | LI-COR | Cat#926-32210 |
| IRDy 800CW Goat anti-Rabbit IgG Secondary Antibody | LI-COR | Cat#926-32211 |

### Supplementary Figures and legends

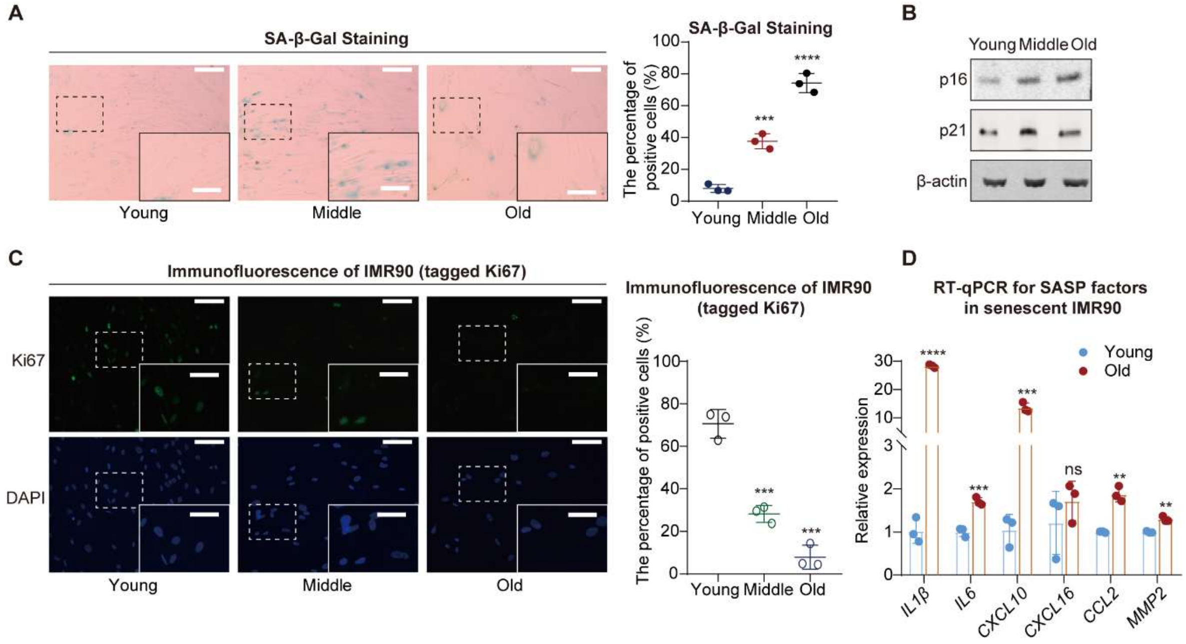

**Figure S1 The expression of various senescence markers during IMR90 cell senescence.** (**A**) SA-β-Gal staining (left) and statistical analysis of SA-β-Gal ratios (right) during IMR90 cell senescence. Scale bars, 200 μm. Enlarged scale bars, 100 μm. (**B**) Western blot analysis showing the protein levels of p16 and p21 in IMR90 cells at different passage. (**C**) Ki67 immunofluorescence experiment (left) and statistical analysis of Ki67 intensity (right) during IMR90 cell senescence. Scale bars, 80 μm. Enlarged scale bars, 40 μm. (**D**) RNA expression of SASP factors during IMR90 cell senescence, with blue bars representing the young group and red bars representing the old group. A two-tailed t-test was employed, ns indicates no significant difference, **$p < 0.01$, ***$p < 0.001$, ****$p < 0.0001$.

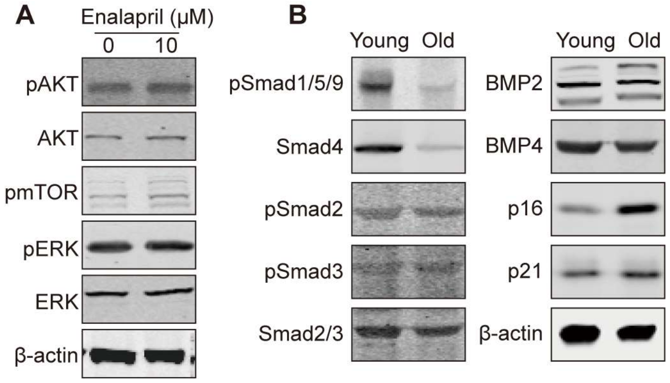

**Figure S2 Alterations in critical regulators of cellular senescence.** (**A**) Western blot analysis showing the changes in protein levels of key targets in classical senescence signaling pathways following enalapril treatment. (**B**) Western blot analysis showing the changes in Smad and BMP-related protein levels during IMR90 cell senescence.

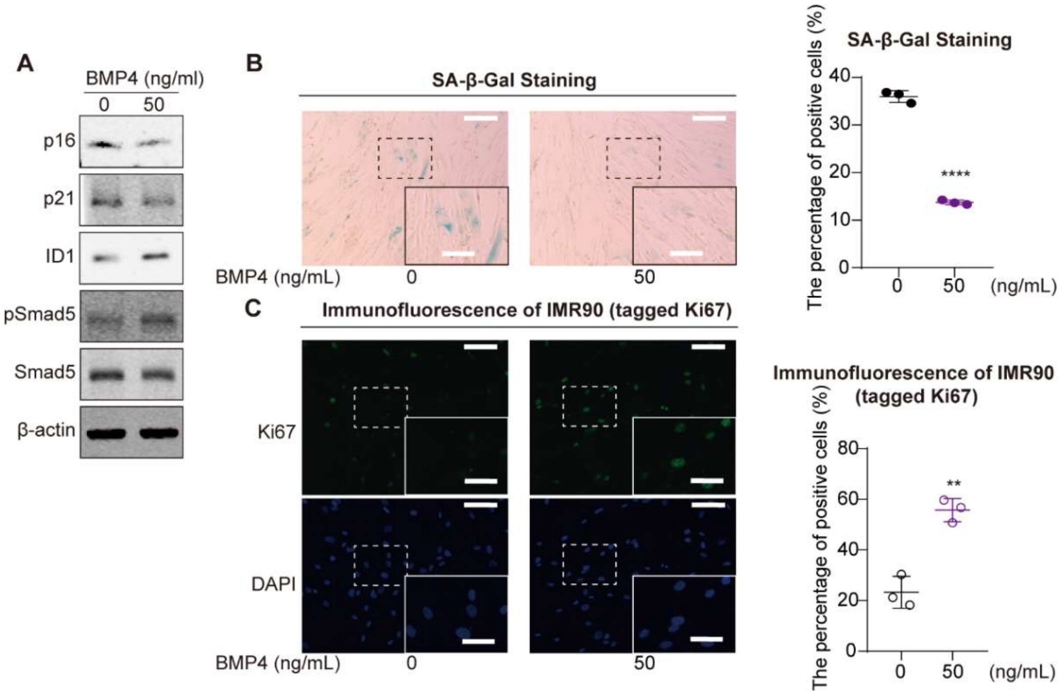

**Figure S3 pSmad1/5/9 alleviates cellular senescence phenotypes.** (**A**) Western blot analysis showing the changes in protein levels following BMP4 treatment. (**B**) SA-β-Gal staining (left) and corresponding ratio analysis chart (right) following BMP4 treatment. Scale bars, 200 μm. Enlarged scale bars, 100 μm. (**C**) Ki67 immunofluorescence (right) and corresponding ratio analysis chart (left) following BMP4 treatment. Scale bars, 80 μm. Enlarged scale bars, 40 μm. A two-tailed t-test was employed, $^{**}p < 0.01$, $^{****}p < 0.0001$.

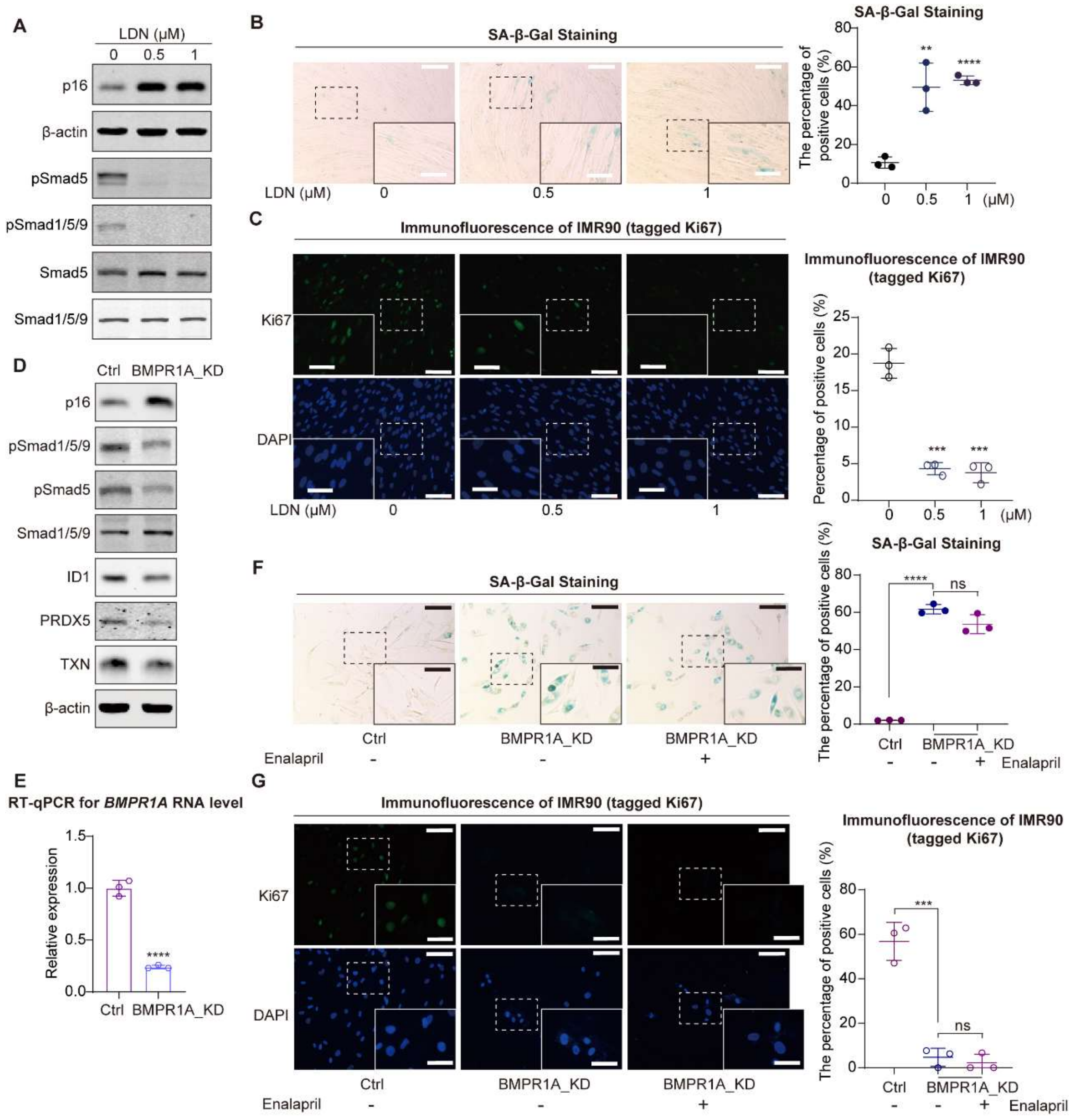

**Figure S4 Reduction of pSmad1/5/9 correlates with cellular senescence.** (**A**) Western blot showing the protein levels after the addition of LDN193189. (**B**) SA-β-gal staining (left) after adding LDN193189 and its corresponding staining proportion statistical chart (right). Scale bars, 200 μm. Enlarged scale bars, 100 μm. (**C**) Immunofluorescence staining for Ki67 (left) after adding LDN193189 along with the corresponding intensity statistical chart (right). Scale bars, 80 μm. Enlarged scale bars, 40 μm. (**D**) Western blot showing the protein levels following BMPR1A knockdown. (**E**) Relative RNA levels of BMPR1A following BMPR1A knockdown. (**F**) SA-β-gal staining (left) and its corresponding staining proportion statistical chart (right) in the control group (Ctrl) and *BMPR1A*-knockdown groups (BMPR1A_KD) with enalapril treatment. Scale bars, 200 μm. Enlarged scale bars, 100 μm. (**G**) Ki67 immunofluorescence (left) and its corresponding staining proportion statistical chart (right) in the control group (Ctrl) and *BMPR1A*-knockdown groups (BMPR1A_KD) with enalapril treatment. Scale bars, 80 μm. Enlarged scale bars, 40 μm. A two-tailed t-test was employed, ns indicates no significant difference, $^{**}p < 0.01$, $^{***}p < 0.001$, $^{****}p < 0.0001$.

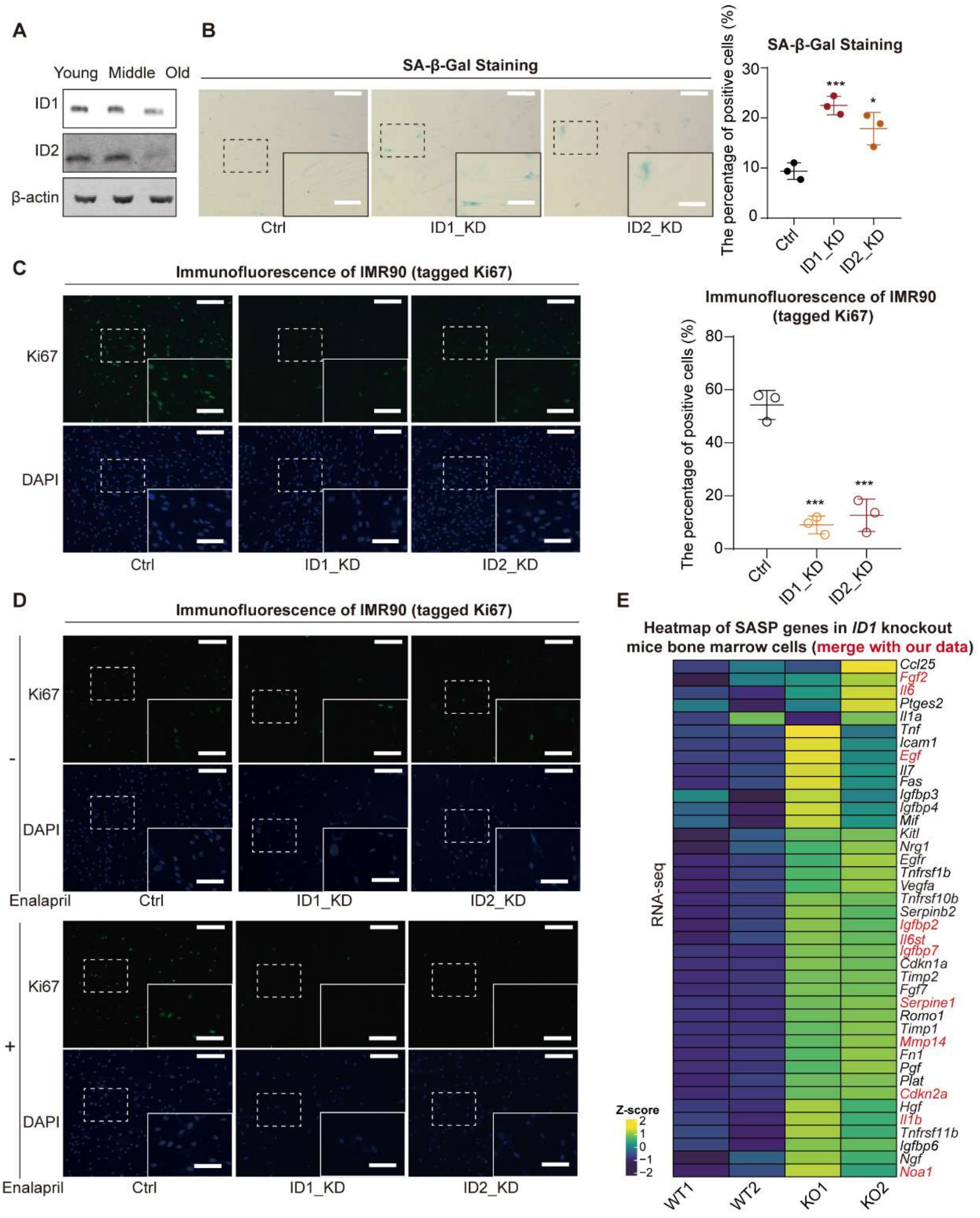

**Figure S5 Knockdown of *ID* accelerates cellular senescence.** (**A**) Western blot showing changes in ID protein levels during cellular senescence. (**B**) SA-β-Gal staining (left) following knockdown of *ID1* and *ID2*, along with corresponding statistical chart (right). Scale bars, 200 μm. Enlarged scale bars, 100 μm. (**C**) Ki67 immunofluorescence (left) following knockdown of *ID1* and *ID2*, along with corresponding statistical chart (right). Scale bars, 80 μm. Enlarged scale bars, 40 μm. A two-tailed t-test was employed, *$p < 0.05$, ***$p < 0.001$. (**D**) Ki67 immunofluorescence in control group (Ctrl) and *ID* knockdown groups (ID1_KD, ID2_KD), with or without enalapril treatment. Scale bars, 80 μm. Enlarged scale bars, 40 μm. (**E**) Heatmap of SASP factor expression in mouse bone marrow cells following *ID* knockout from previously reported data(Fei et al., 2023).

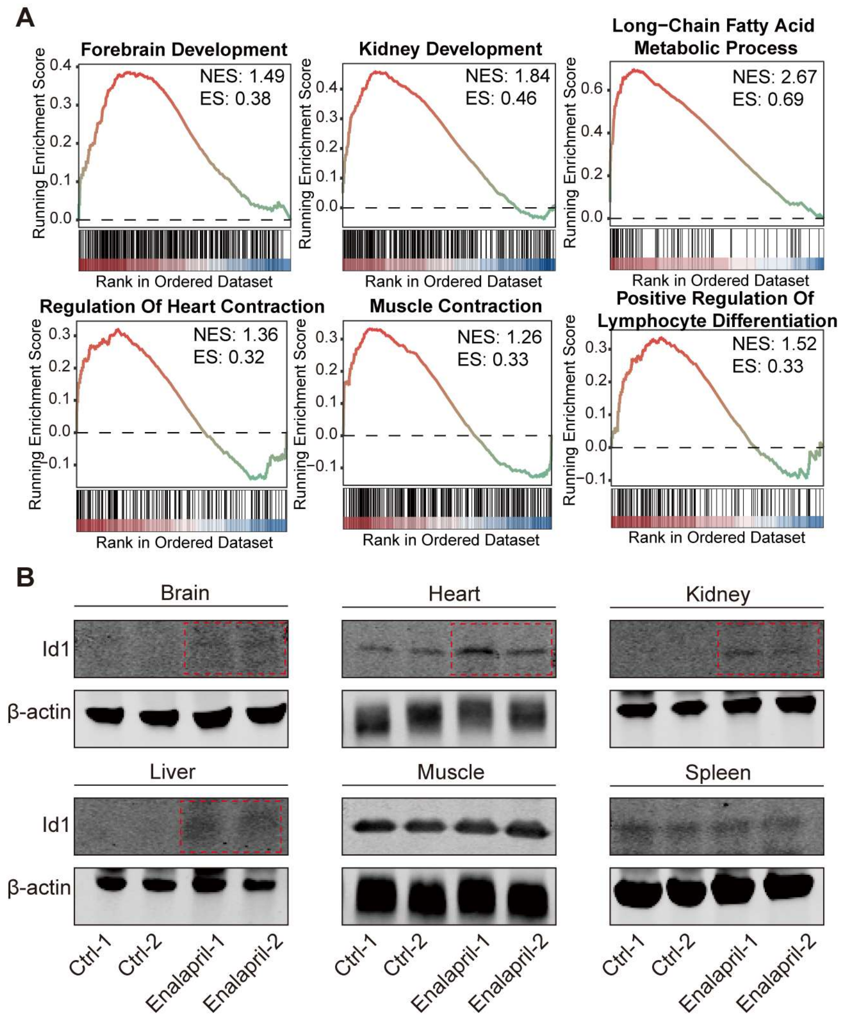

**Figure S6 Effects of enalapril on the expression changes of genes related to organ physiological functions in mice.** (**A**) In multiple organs, enalapril increases the enrichment of GSEA pathways associated with improved physiological function. (**B**)

Western blot analysis showing changes in Id1 protein levels in various organs of mice.

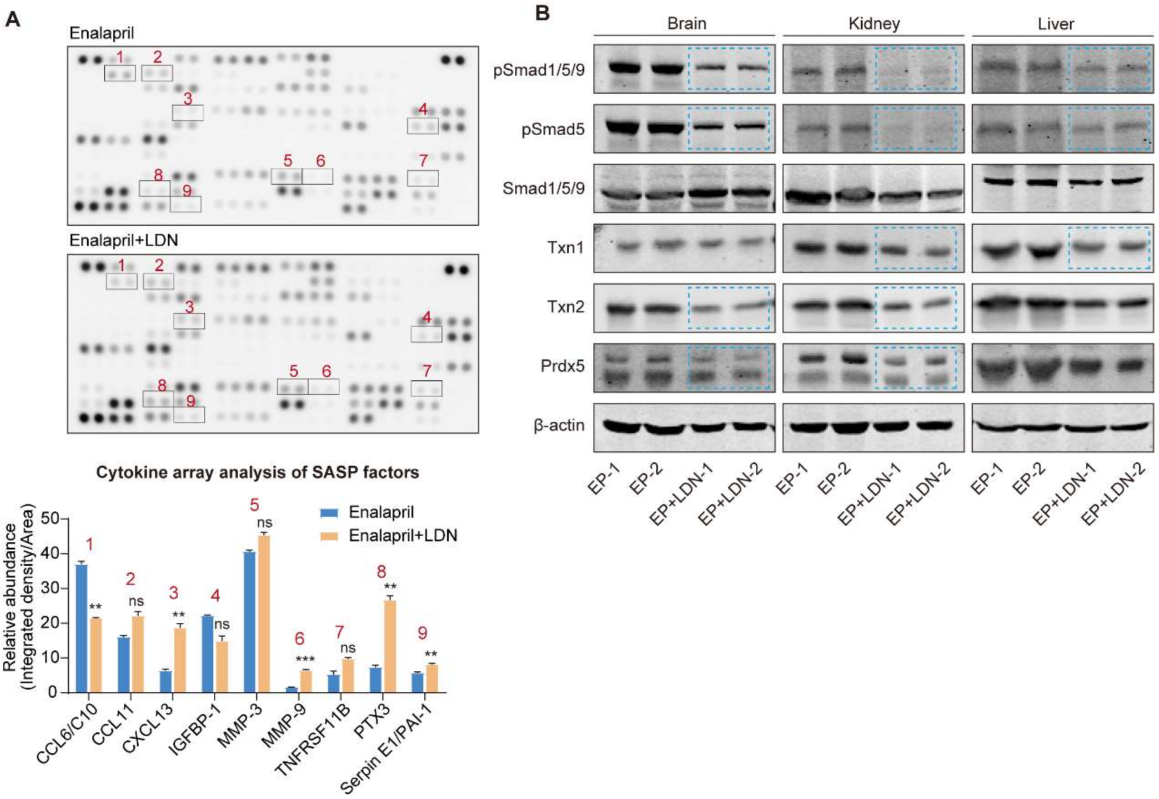

**Figure S7 Effects of reduced pSmad1/5/9 levels on the expression changes of genes in mice.** (**A**) Cytokine array analysis of secreted proteins (above) and relative quantitation (below) of SASP factors in enalapril-treated serum (Enalapril) and co-treated serum (Enalapril+LDN). (**B**) Western blot analysis showing the protein levels of pSmad1/5/9 and antioxidative genes in various organs following enalapril and LDN co-treatment.

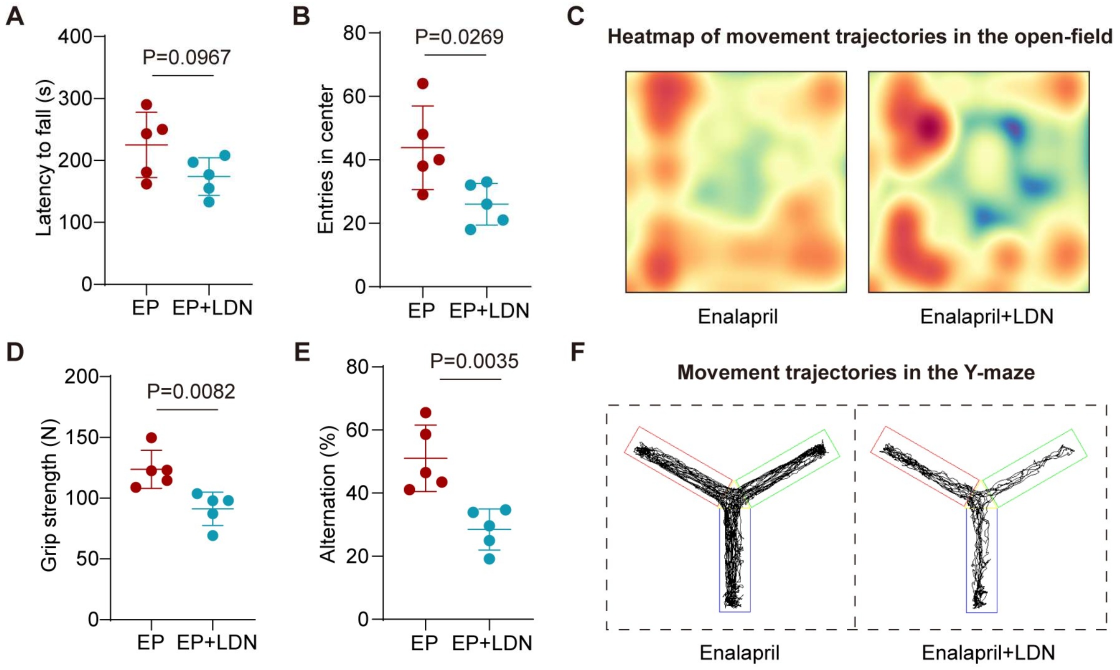

**Figure S8 Effects of reduced pSmad1/5/9 levels on aging-related behaviors in mice.** (**A, D**) Changes in the rotarod fall time (**A**) and grip strength (**D**) of the mice after enalapril and LDN treatment. (**B, C**) The number of entries into the central area (**B**) and heatmap of movement trajectories (**C**) in the open-field test after enalapril and LDN treatment. (**E, F**) Spontaneous alternation rate (**E**) and movement trajectory (**C**) in the Y-maze of the mice after enalapril and LDN treatment.

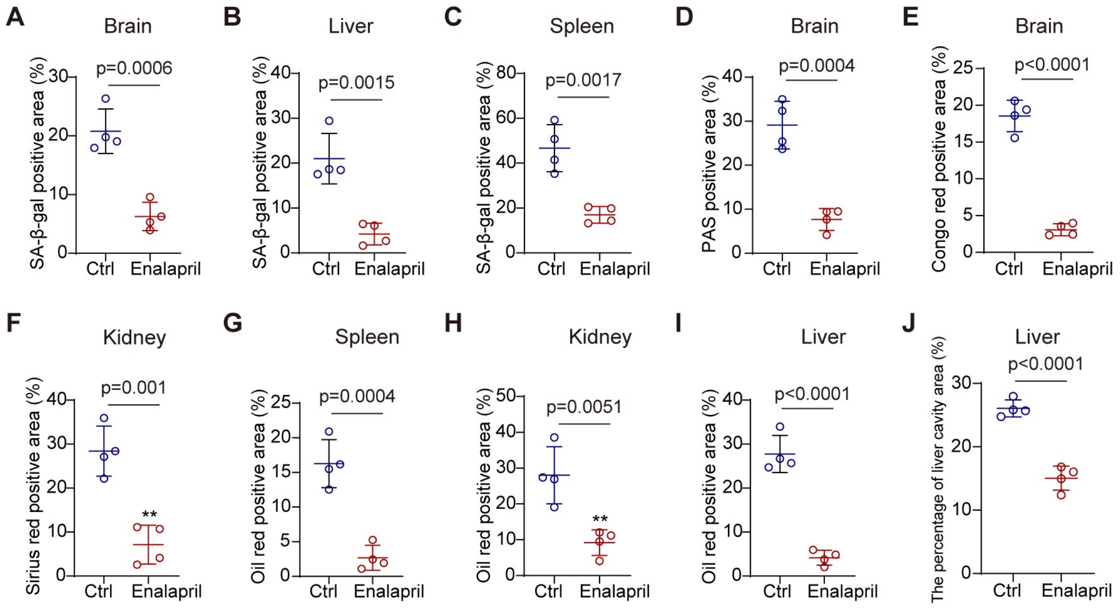

**Figure S9 Enalapril ameliorates pathological phenotypes in aged mice.** (**A-C**) SA-β-Gal staining quantification chart of the brain (**A**), liver (**B**) and spleen (**C**) after enalapril treatment. (**D, E**) Periodic Acid-Schiff (PAS) staining (**D**) and Congo red staining (**E**) quantification chart of the brain after enalapril treatment. (**F**) Sirius red staining quantification chart of the kidney after enalapril treatment. (**G-I**) Oil Red O staining quantification chart of the spleen (**G**), kidney (**H**) and liver (**I**) after enalapril treatment. (**J**) Hematoxylin and eosin (H&E) staining quantification chart of the liver after enalapril treatment.
